# Myosin Orientation in a Muscle Fiber Determined with High Angular Resolution Using Bifunctional Spin Labels

**DOI:** 10.1101/513556

**Authors:** Yahor Savich, Benjamin P. Binder, Andrew R. Thompson, David D. Thomas

## Abstract

We have measured the orientation of the myosin light chain domain (lever arm) elements in demembranated muscle fibers by electron paramagnetic resonance (EPR), using a bifunctional spin label (BSL), with angular resolution of 4 degrees. Despite advances in X-ray crystallography and cryo-electron microscopy (cryo-EM), and fluorescence polarization, none of these techniques provide high-resolution structural information about the myosin light chain domain under ambient conditions in a muscle fiber. Two cysteines, 4 residues apart, were engineered on two α-helices in the myosin regulatory light chain (RLC), permitting stereoselective site-directed labeling with BSL. One labeled helix (helix E) is adjacent to the myosin lever arm, the other helix (helix B) is located farther apart from the motor domain beyond the “hinge” of the myosin. By exchanging BSL-labeled RLC onto oriented muscle fibers, we obtained EPR spectra that determined angular distributions of BSL with high resolution, which enabled the accurate determination of helix orientation of individual structural elements with respect to the muscle fiber axis. In the absence of ATP (rigor), each of the two labeled helices exhibited both ordered (σ ~ 9-11 degrees) and disordered (σ > 38 degrees) populations. We used these angles to determine the orientation of the myosin lever arm, concluding that the oriented population has lever arms that are perpendicular to the muscle fiber axis. This orientation is ~33 degrees different than predicted from a standard “lever arm down” model based on cryo-EM of actin decorated with isolated myosin heads, but it is compatible with fluorescence polarization and EM data obtained from muscle fibers. The addition of ATP, in the absence of Ca^2+^, shifted the orientation to a much more disordered distribution.

**Summary:** We used electron paramagnetic resonance to determine the orientation of elements within the myosin regulatory light chain in skinned skeletal muscle fibers. A bifunctional spin label provided sufficient resolution to detect an ordered population of lever arms perpendicular to actin.

## INTRODUCTION

Muscle contraction is a process in which the nanometer-sized force-bearing elements of myosin contribute to macroscopic contraction of the tissue. Despite recent technical advances in cryo-EM and X-ray crystallography, important structural details of the mechanism of force generation by myosin remain unclear. The essential light chain (ELC) and RLC bind to an α-helix on the myosin heavy chain, forming the light-chain domain, which acts as a lever arm for producing force and movement. Either due to structural flexibility of this complex, or to sample inhomogeneity, the above methods have not definitively determined the structure of the lever arm, even in the simplest biochemical state – rigor (no ATP). In addition, none of the above-mentioned structural techniques provide structural information under physiological conditions (not frozen or crystallized) in muscle fibers. Fluorescence polarization anisotropy does not have this limitation, and has been used to provide orientational information about the ELC (Knowles et al., 2008) and RLC (Brack et al., 2004; Romano et al., 2012; Fusi et al., 2015) in skinned fibers. However, fluorescence has low angular resolution, so this approach requires data from multiple labeling sites and complex data analysis.

Electron paramagnetic resonance (EPR) of nitroxide spin labels offers high orientational resolution (1–5 degrees) under physiological conditions, which is needed to overcome the above limitations (Thomas and Cooke, 1980; Cooke et al., 1982; Fajer et al., 1986; Fajer et al., 1988; Arata, 1990; Hambly et al., 1991, 1992; Zhao et al., 1996; Thomas et al., 2009; Nogara et al., 2016). The sensitivity of the EPR spectrum to the angle between the nitroxide’s π orbital (*z*_*N*_-axis in Fig. 1B) and the magnetic field 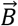 light blue arrow in Fig. 1B), and to the rate and amplitude of rotational motion, provides high-resolution information about the orientation and rotational motion of a spin label relative to the magnetic field. Use of a deuterated spin label increases the angular resolution of EPR (Fajer, 1994b, a), but monofunctional (flexible) attachment of the label produces ambiguity concerning orientation of the protein structural elements. BSL (Fig. 1) provides rigid and stereospecific attachment of the probe to an α-helix (Wilcox et al., 1990; Fleissner et al., 2011; Sahu et al., 2017), thus providing unambiguous information about helix orientation (with ~1 degree resolution) and rotational motion, as demonstrated previously in applications to actin-bound myosin S1 (Arata et al., 2003; Rayes et al., 2011; Binder et al., 2015; Thompson et al., 2015; Binder et al., 2018). Due to increasing interest in thick filament muscle regulation, it is important to study both the actin-attached and -detached states of myosin with BSL’s high angular resolution, given that the orientation of the lever arm in multiple biochemical states in contracting muscle remains ambiguous (Irving, 2017).

**Fig. 1.**
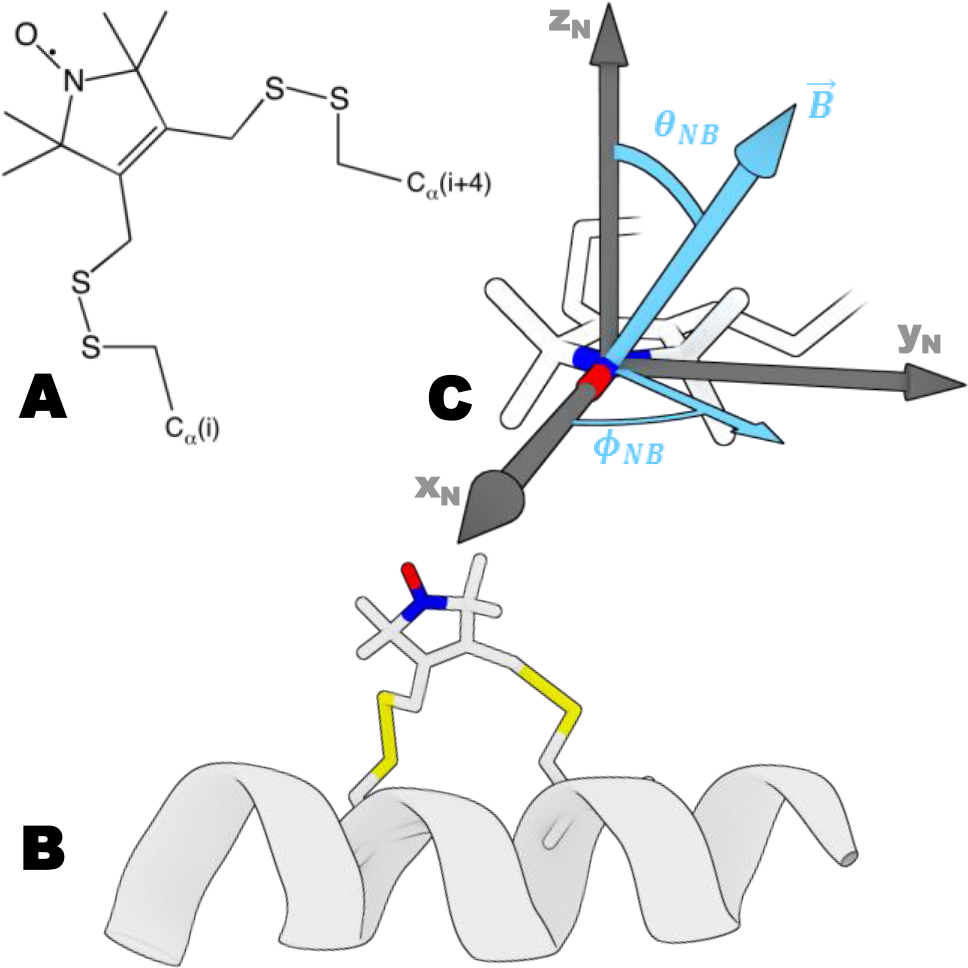
(A) Chemical structure of BSL reacted with two Cys residues. (B) BSL bound stereospecifically to an α-helix at positions i and i+4. (C) Angles that define the orientation of the nitroxide (defined by axes x_N_, y_N_, z_N_) relative to the applied magnetic field 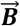 these angles directly determine the high-resolution orientation dependence of the EPR spectrum. Coordinates of the BSL are taken from (Binder et al., 2018).

In the present study, we exchanged BSL-labeled RLC onto the myosin lever arm in permeabilized rabbit muscle fiber bundles, and measured the orientation of helices in the N- and C-lobes in the absence of nucleotide (rigor) and in the presence of ATP (relaxation).

## METHODS

### Protein and muscle fiber preparations

Wild type rabbit skeletal regulatory light chain (RLC) insert (UniProtKB entry: MLRS_RABIT; P02608) was obtained from Genscript. Wild type chicken gizzard smooth muscle RLC (smRLC) insert was as in (Mello and Thomas, 2012). Mutants of skRLC and smRLC were obtained by Q5 site-directed mutagenesis kit (New England BioLabs) in pET3a vector. Constructs were verified via sequencing at University of Minnesota Genomics Center. Purified plasmid (ZymoPURE Plasmid Miniprep kit) was transformed into BL-21AI strain of E. coli. Protein was expressed and prepared via inclusion bodies purification as in (Nelson et al., 2005). Endogenous cysteines were replaced by alanines, and a di-cysteine BSL labeling motif was introduced for each construct on helix B (D53C-A57C) and helix E (G103C-V107C) in skRLC, and (D56C-S60C) in smRLC. Skinned rabbit psoas muscle fiber bundles were dissected, permeabilized and stored in fiber storage buffer (Prochniewicz et al., 2008) with 4mM DTT added. Heavy meromyosin (HMM) was purified from rabbit skeletal muscle (Muretta et al., 2015).

### Preparation of BSL-RLC

Di-Cys RLC mutants in labeling buffer (30mM Tris, 50mM KCl, 3mM MgCl_2_, pH 7.5) were incubated at 4°C with 5mM DTT for 1hr to ensure reduction of engineered cysteine residues prior to labeling. DTT was removed using Zeba Spin desalting columns (Thermo Scientific), and RLC was incubated at 4°C for 1hr in 5-fold molar excess of bifunctional 3,4-Bis-(methanethiosulfonylmethyl)-2,2,5,5-tetramethyl-2,5-dihydro-1H-pyrrol-1-yloxy spin label (BSL; Toronto Research Chemicals). The covalent disulfide double bond between BSL and the 𝛼-helix backbone produces bifunctional stereospecific attachment in this labeling conditions (Fig. S2 of Binder et al., 2015). Following incubation, excess spin label was removed and the protein was exchanged into rigor buffer (25 mM Imidazole, 10 mM EGTA, 100.3 mM KPr, 1.5 MgAc_2_, pH 7.1) using Zeba Spin desalting columns. Labeling efficiency (spin labels attached per protein) was determined to be >80% based on double integration of EPR spectra. Mass spectrometry (performed on Agilent 7200B Quadrupole Time-of-Flight GC/MS) shifted unlabeled populations by 228 Da (SD = 2 Da, n = 5), as predicted for labels molecular weight (230 Da) attachment.

### BSL-RLC exchange onto HMM and decoration of fibers with BSL-RLC-HMM

BSL-labeled RLC was exchanged onto HMM (Muretta et al., 2015) by combining the two proteins (3:1 RLC:HMM, mol/mol) in 50 mM Tris pH 7.5, 120 mM KCl, 12 mM EDTA. Samples were incubated for 10 minutes at 30°C, followed by addition of 12 mM MgCl_2_ and incubation on ice for 15 min. Free RLC was subsequently removed using Amicon Ultra Centrifugal Filters, 0.5 mL, 50K MWCO (EMD Millipore), and finally samples were exchanged into rigor buffer. Actin in fiber bundles was decorated with the resulting BSL-RLC-HMM complex by circulating protein samples through a 25 μL glass capillary (Drummond Scientific) containing a tied fiber bundle.

### BSL-RLC exchange into permeabilized fibers

Glycerinated rabbit psoas fibers were dissected into bundles, ranging from 0.3-0.5 mm in diameter and 3-5 cm in length, and were tied at each end using surgical silk. The tied fiber bundles were then held in place at a fixed length within a 25 μL glass capillary (Drummond Scientific), with the ends of the sutures affixed to the capillary using short sections of silicone tubing. Buffer exchange was accomplished with a Masterflex C/L peristaltic pump (Cole-Parmer) at the flow rate of 0.5 mL/min. Exchange of BSL-labeled RLC constructs onto permeabilized fibers was done as described previously (Mello and Thomas, 2012), with the exception that the concentration of DTT in the wash before the acquisition was 0.5 mM instead of 30 mM. TnC was reconstituted during one hour incubation at 4°C. Rabbit skeletal TnC was purchased from Life Diagnostics. It was shown previously that this exchange and reconstitution protocol is complete (> 90% reconstitution for both RLC and TnC) and has no significant effect on function, as measured by Ca-dependent myofibrillar MgATPase assays (Mello and Thomas, 2012). Rigor and relaxation solutions used during acquisition were as described in (Fusi et al., 2015). Ionic strength and pH during acquisition were maintained at 150 mM and 7.1, respectively.

### EPR spectroscopy

EPR was performed on a Bruker EleXsys E500 X-band spectrometer (9.6 GHz). Spectra of BSL-RLC and BSL-RLC-HMM were acquired in an ER4122 SHQ spherical resonator. Parallel and perpendicular field experiments on fiber bundles were performed in TM_110_ 4103TMA and TE_102_ 4104OR-R resonators, respectively. Minced fiber measurements were performed in a quartz flat cell in the TE_102_ 4104OR-R cavity. Sample temperature was set to 4oC in all experiments. Sweep width was 120 G with 1024 points per spectrum, but only 100 G is displayed in figures. The center magnetic field (*B*_*c*_) was set according to *B*_*c*_ =ν/2.803 *MH𝑧/𝐺*. Conversion time and time constant were 20.48 ms. Modulation amplitude and microwave power were 1-2 G and 2-20 mW, depending on the cavity.

### Data analysis

EPR spectra were background-subtracted, normalized by dividing by the second integral, and analyzed as described previously to determine the angular distribution of spin labels relative to the muscle fiber axis, assuming the rigid limit regime due to the immobilization granted by the BSL spin label (Binder et al., 2015). Magnetic tensor values (g and T, defining the orientational dependence of the spectrum) were determined from completely disordered samples (minced fibers). The angular distribution was fit to Gaussian functions for both angles *θ*_*NB*_ and *ϕ*_*NB*_, (Fig. 1C), with standard deviations *σ_θ_* and *σ_ϕ_*. Sensitivity of EPR to *θ*_*NB*_ in systems with axial symmetry (as in oriented muscle fibers) is considerably greater than the sensitivity to *ϕ*_*NB*_, so we focus on *θ*_*NB*_ values for molecular modeling. Confidence intervals were determined from the cumulative distribution function, comparing the ratio of the residual sum of squares of the fits with the residual sum of squares of the best fit.

### Molecular modeling

Angle measurements derived from atomistic models were calculated using custom extensions written for Visual Molecular Dynamics 1.9.3 (VMD) (Humphrey et al., 1996). Minimization of models was performed using SciPy and NumPy packages of Python. Structure images were rendered using Blender, Version 2.79 (Blender Foundation).

### Supplementary material summary

Table S1 summarizes orientational parameters snown on the Fig. S1. Fig. S2 shows the cumulative distribution function of the F-ratio distribution for the key parameters in the present paper. Fig. S3 indicates the effect of the AMPPNP on the oriented BSL-RLC-fiber labeled on the B helix.

## RESULTS

### Detecting anisotropy of BSL-RLC-fiber and motion BSL-RLC-HMM in EPR

We performed three types of EPR experiments on muscle fiber bundles: with the muscle fiber axes (1) parallel or (2) perpendicular to the applied magnetic field, or (3) randomly oriented due to mincing, thus eliminating anisotropy. The parallel spectrum line shape (Fig. 2A, magenta) typically shows the greatest difference from that of the minced fiber (Fig. 2A, green). The perpendicular experiment is also sensitive to orientation (Fig. 2A, orange), but apparent disorder is caused by the helical symmetry of the actomyosin complex about the muscle fiber axis. Therefore, we focus on the parallel spectra, compared with spectra of minced fibers.

**Fig. 2.**
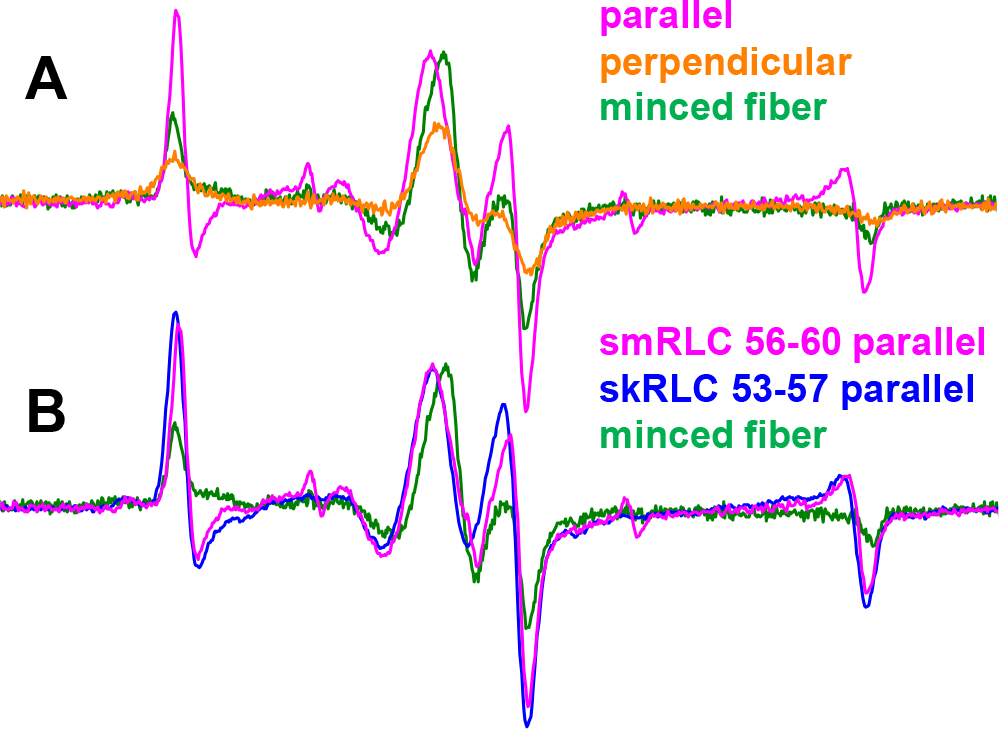
**(A)** The effect of changing fiber orientation on EPR spectrum for smRLC helix B (56-60). **(B)** The parallel field experiment on helix B of skRLC and smRLC homologs labeled at equivalent sites.

Our previous work on myosin in skeletal fiber bundles and HMM involved smooth muscle RLC (smRLC) from chicken gizzard (Mello and Thomas, 2012; Muretta et al., 2015). Here we extend this to the skRLC homolog. Fig. 2B shows that *θ*_*NB*_ orientations of smRLC and skRLC are hardly distinguishable, which is consistent with fluorescence polarization experiments (Romano et al., 2012). Therefore, to maximize the physiological relevance of our work, we focus on skRLC.

We exchanged skeletal RLC constructs on HMM to and measured EPR in solution to understand the motional regime of the RLC and RLC-BSL. In randomly oriented samples (e.g., proteins in solution or minced fibers), the rotational correlation time (characteristic time for diffusive rotation by one radian) of the spin label was calculated from 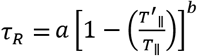 where 2*T*^′^_∥_ is the splitting between the low- and high-field features of the observed spectrum (Fig. 3), 2*T*_∥_ is the rigid-limit splitting (69-71 G, depending on local polarity), with *a* = 5.4 × 10^−10^*s* and *b* = −1.36 (Goldman et al., 1972). Exchanging BSL-RLC (MW 19,000) onto HMM (MW 350,000) leads to 2*T*^′^_∥_ = 70.6 G (SD = 0.2 G, n = 4) (Fig. 3, black), indicating rigid immobilization of the probe on the ns time scale. We conclude that BSL is rigidly immobilized when bifunctionally attached to RLC. This is consistent with the rigid and stereospecific immobilization seen previously with BSL attached to other proteins (Her et al., 2018).

**Fig. 3.**
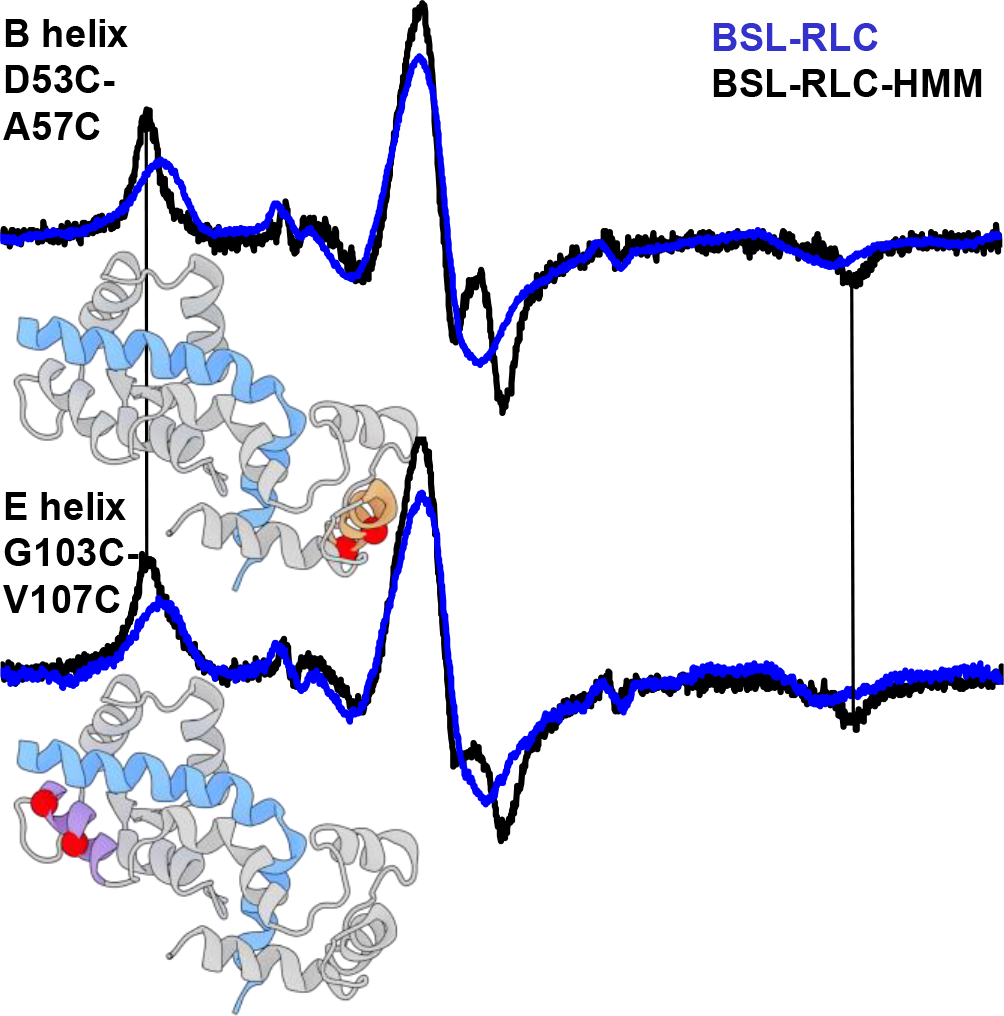
EPR spectra of BSL-RLC (blue) and BSL-RLC-HMM (black) in solution (randomly oriented). Structural models show RLC in grey and myosin heavy chain in blue. The position of the label is depicted by red spheres on the B helix (orange) of the N-lobe and the E helix (magenta) of the C-lobe (PDB: 5H53, Fujii and Namba, 2017). Vertical bars indicate the positions of outer peaks of the HMM spectra, which indicate a lack of sub-microsecond rotational motion. Field sweep is 100 G.

### Orientation of the probes in fiber bundles

Following the RLC-fiber exchange protocol, we performed EPR measurements on helices B and E. (Fig. 4 Fig. 5). With a single Gaussian oriented component, the B helix in the N-lobe was fit by (*θ*_*NB,B*_, σ_*θ,B*_) ≈ (4 ± 4°, 9° ± 3°) and the E helix in the C-lobe was fit by (*θ*_*NB,E*_, σ_*θ,E*_) ≈ (81 ± 4°, 11 ± 3°) (Fig. 4C, blue). Confidence intervals were determined from the cumulative distribution function of the F-ratio distribution, comparing the ratio of the residual sum of squares of the fits with the residual sum of squares of the best fit. The intervals of 95% are depicted by horizontal lines (Fig. S2).

**Fig. 4.**
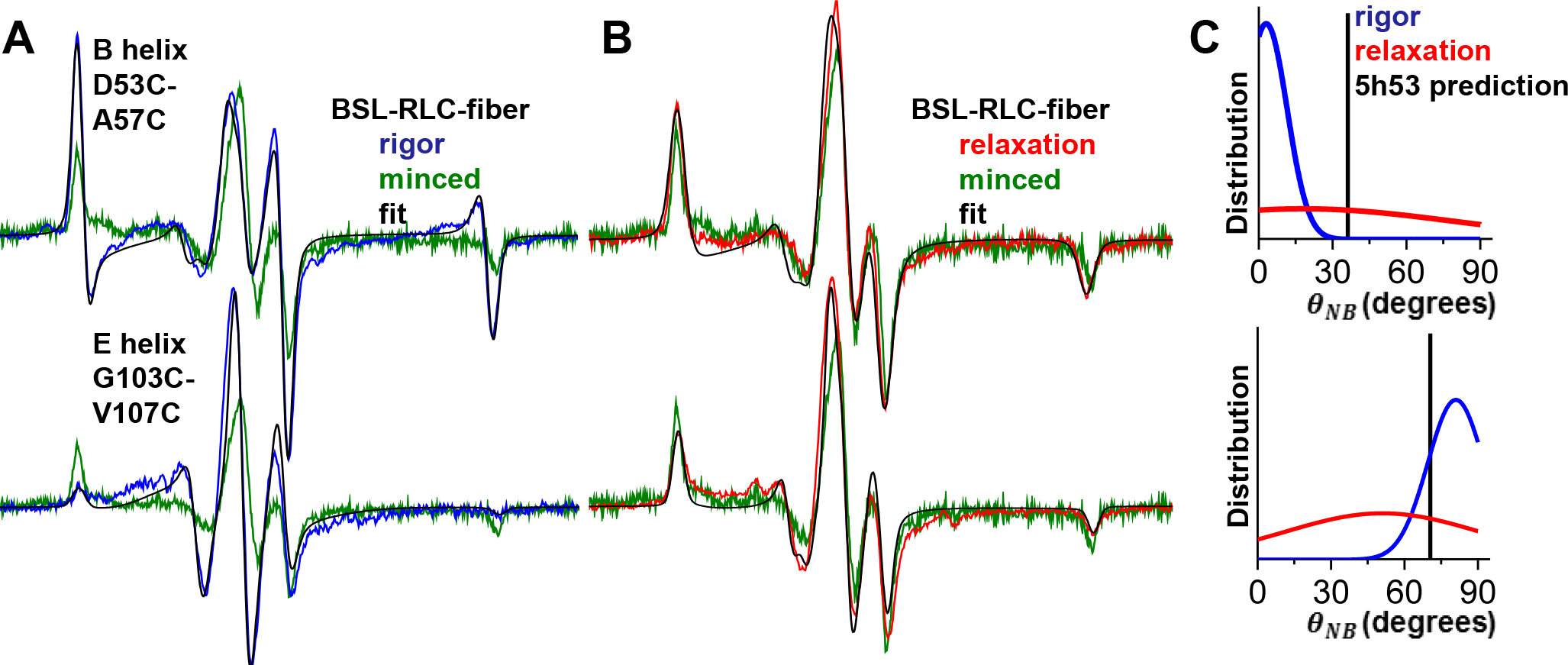
EPR of BSL-RLC-fiber in **(A)** rigor (blue) and **(B)** relaxation (red). Fits are in black and the minced fiber data (isotropic control) is green. Field sweep is 100 G. The *θ*_*NA*_ distribution in **(C)** for nucleotide-free (blue) and ATP-bound (red) state, derived from corresponding helices. Vertical bars represent predicted angles from the (PDB: 5H53) model.

**Fig. 5.**
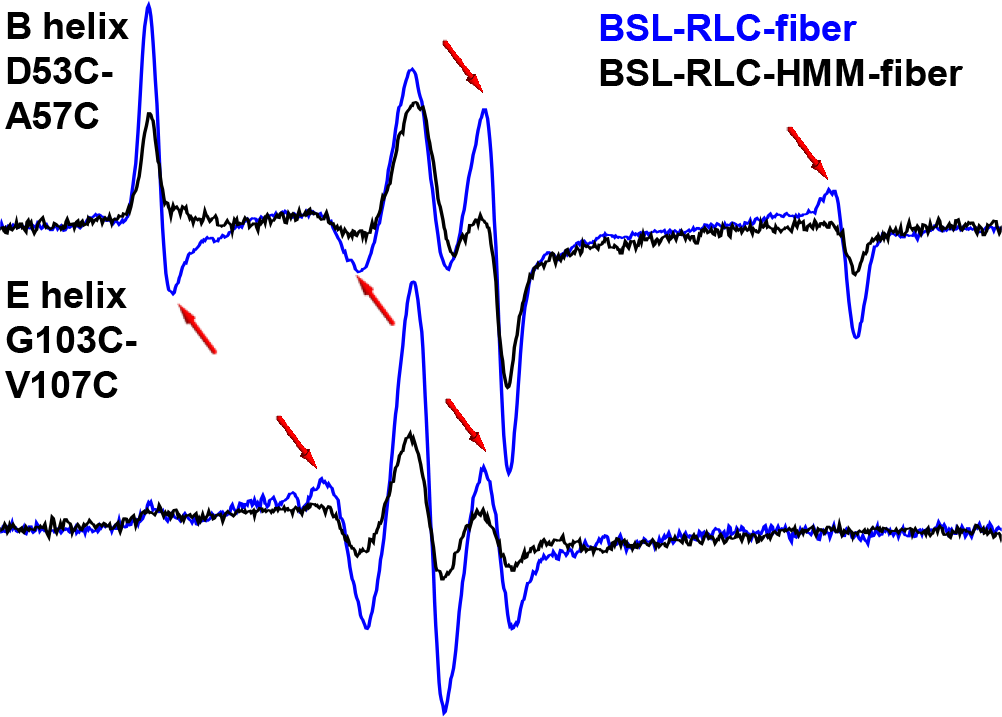
EPR spectra of BSL-RLC-HMM-fiber (black) and BSL-RLC-fiber (blue) in parallel orientation. Red arrows indicate the most prominent spectral features of the oriented component. Field sweep is 100 G.

Addition of ATP, inducing relaxation, produces spectra that are very similar to those of minced fibers (compare red and green in Fig. 4B). Fitting these spectra with a single Gaussian component resulted in σ_*θ,B*_ ≈ 60° and σ_*θ,E*_ ≈ 40°. This degree of disorder is consistent with RLC-RLC distance measurements performed by double electron-electron resonance (DEER) on smooth muscle myosin filaments labeled with a monofunctional spin label (Vileno et al., 2011), where it was observed that probes on the N-lobe exhibited greater disorder than on the C-lobe. Previous fluorescence polarization experiments also show substantial orientational disorder in relaxation at 4°C (Fusi et al., 2015), consistent with our results.

### Probes are much less oriented in BSL-RLC-HMM-decorated fibers

To ensure that the measured orientation in a fiber bundle stems from RLC stereospecifically bound to the IQ domain of the actin-attached myosin, we exchanged BSL-RLC onto HMM, then decorated a fiber bundle with BSL-RLC-HMM. The spectrum of the E helix in this experiment is similar to that of the BSL-RLC-fiber (Fig. 5, bottom, Table S1), except that the mole fraction of the oriented component is less (about half). The spectrum of the B helix is also less ordered (compare blue and black, Fig. 5, top; Table S1). In both cases, disorder increases when myosin is cut at the HMM/LMM junction. The greater disorder of the B helix is consistent with its more distal location (compared with the E helix) on the lever arm, near the flexible hinge. For both the E and B helices, the orientation of BSL-RLC is greater for the labeled fiber bundle (Fig. 5; Table S1) than for HMM. We conclude that BSL-RLC labeling of the lever arm must be at least as specific in intact fibers as in HMM-decorated fibers.

Previously published FRET experiments, with a donor on the E helix of RLC bound to HMM and an acceptor bound to the active site, produced a well-defined donor-acceptor distance, implying stereospecificity. Biochemically-initiated kinetic steps in the ATPase cycle, with labeling sites on exchanged RLC and the catalytic domain, lead to 25-30% FRET changes during ATP cycling, consistent with a stereospecific population of RLC bound during lever arm transitions (Muretta et al., 2015; Rohde et al., 2017). Non-specific binding of RLC to demembranated fibers can obscure the measurement of orientation or lead to an inconclusive data interpretation. The RLC-fiber exchange protocol that was used in the present study yields <15% of non-specifically bound RLC (Mello and Thomas, 2012).

In order to interpret these results in terms of the movement of the lever arm, we performed molecular modeling of the actin-attached state in the absence of nucleotide, combining the orientational distributions from both B and E helices (Fig. 4C, blue).

### Model of a lever arm in a nucleotide-free state

Having determined orientational distributions of BSL on the B and E helices of the RLC, we sought to compare those measurements with established atomic models of the actomyosin complex. We began with a recent cryo-EM structure derived from rabbit skeletal muscle (PDB: 5H53) (Fujii and Namba, 2017) which used the squid 3I5G (Yang et al., 2007) crystal structure for docking and refinement of myosin coordinates. Coordinates for BSL were obtained from a crystal structure of the label bound to a helix in T4 lysozyme (Fleissner et al., 2011). The presence of an unresolved EM density in the above-mentioned reference suggests a small alternative conformation of BSL. We used a conformation of BSL that was obtained from global analysis of orientational EPR data and double electron-electron resonance data (Binder et al., 2018). BSL was modeled onto both experimental RLC sites by alignment of backbone atoms, with the orientation of each label adjusted to reflect conformational refinements from recent EPR studies (Binder et al., 2015). A *θ*_*NB*_ value was subsequently calculated from the model for each label, with the model’s actin filament axis assumed to be parallel to the external magnetic field (as it would be in a parallel-oriented fiber experiment). Initial results gave poor agreement with the orientational distribution centers derived from our EPR data: *θ*_*NB*_ for helix B was 37.1° (difference of +33.1°), and *θ*_*NB*_ for helix E was 70.4° (difference of −10.6°) (Fig. 5C, blue).

This initial result was not surprising, as (1) the myosin captured in 5H53 structure was not resolved in the lever arm domain, and (2) the structure is of frozen myosin S1, and thus the lever arm, might have a very different orientation than measured in our EPR experiments with HMM and full-length myosin in an *in situ* demembranated fiber bundle. Therefore, we sought to find the orientation of the myosin lever arm that best accommodates the measurements derived from EPR.

We began by aligning additional myosin S1 structures to our actomyosin model, choosing several structures solved in various biochemical states with different lever arm orientations (Houdusse et al., 2000; Himmel et al., 2002), (Fig. 6). Next, we used an established method to calculate a vector corresponding to the lever arm helix in each structure (Enkhbayar et al., 2008), and subsequently found the approximate center of rotation for the lever arm by calculating the average point of nearest convergence for all vectors (Fig. 6). Next, we applied a series of arbitrary 3D rotations about that defined center to the portion of our model comprising the lever arm, RLC, and attached labels. After each transformation, we calculated new *θ*_*NB*_ values from the model, and assessed their deviation from the corresponding EPR-derived measurements. Minimizing these deviations by least-squares optimization yielded a new orientation of the lever arm that is compatible with our experimental findings (Fig. 6).

**Fig. 6.**
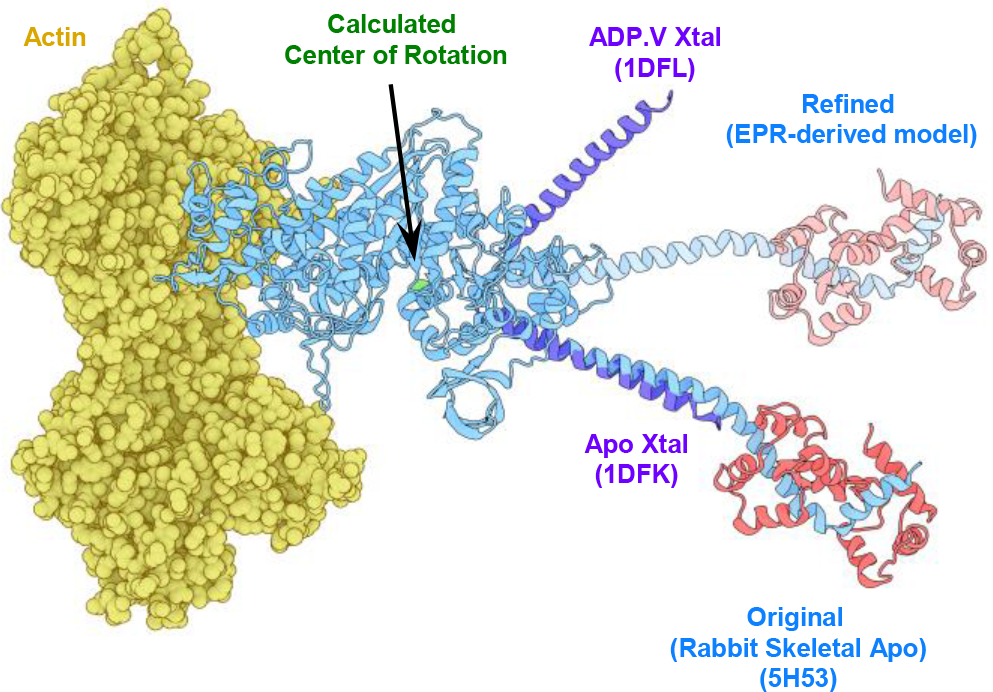
Refinement of the rigor actomyosin model. In blue: the initial model of skeletal myosin attached to actin (yellow) with RLC in red (5H53). Catalytic domains of myosin in other states were aligned with the catalytic domain of 5H53 to find the plane of lever arm rotation (purple) – 1DFL and 1DFK (Houdusse et al., 2000). Center of rotation is denoted by a green sphere. In pale blue: lever arm orientation derived from EPR-derived orientations of helices B and E on the RLC (pale red).

While there are of course many potential 3D rotations that could bring the angles into agreement, the structure depicted in (Fig. 6) represents the solution that renders the smallest possible perturbation on our initial cryo-EM-based model. The optimal rotation relative to the starting structure is parameterized by Euler angles α = 0°, 𝛽 = −33.2°, and γ = +3.3°, in the model’s global coordinate space using the ZYZ-convention (as in the SI of the (Binder et al., 2018)). The small magnitude of both α and γ indicate minimal azimuthal rotation relative to the actin axis, which was oriented along the global z-axis in our model. While the β value indicates a significant alteration in the lever arm tilt relative to actin, it is striking to observe that this EPR-informed solution falls within the same rotational plane as the lever arm’s in other atomic structures of S1 (Fig. 6). We will refer to this rotational plane as the “powerstroke plane” in subsequent discussion. We also applied the above procedure to a structure of chicken skeletal myosin (PDB ID: 2MYS). In the 2MYS model, our labeling sites correspond to E49C-A53C and G99C-V103C. The search for an EPR-compatible rotation of the lever arm converged on a configuration with α = 0°, β = −48.7°, and γ = +20.2°. Thus, not only did the 2MYS-based rotation consist of much more drastic adjustments than were rendered on 5H53, but also placed the lever arm significantly outside of the power-stroke plane. Hence, the smallest structural adjustment of the lever arm accommodating our data is a rotation α = 0°, β = −33.2°, and γ = +3.3°, which brings 5H53 lever arm down to a more perpendicular orientation (Fig. 6).

## DISCUSSION

### Summary of Results

We used BSL to determine the orientation of specific structural elements of skeletal muscle myosin. We studied BSL-labeled fiber bundles in three configurations – parallel to the magnetic field, perpendicular to the magnetic field, and minced (randomly oriented) (Fig. 2A). We found that smRLC, used in previous studies (Mello and Thomas, 2012), reports similar orientation compared to skRLC, consistent with previous results (Romano et al., 2012).

BSL has been previously shown to adopt a highly stereospecific conformation on the α-helices of the myosin motor domain (Binder et al., 2015; Binder et al., 2018). In the present study, we found that BSL was immobilized when bound to helices B and E on the RLC (Fig. 3).

Since our angular predictions for demembranated fiber bundles did not agree with S1 5H53 cryo-EM model data, we found a minimum rotation of the myosin lever arm that brings it to agreement with our angular parameters (Fig. 6). The rotation corresponds to a myosin state with lever arm perpendicular to the fiber axis. A key result of the present study is a method to resolve the oriented components of myosin beyond the motor domain in a muscle fiber under ambient conditions, with high angular resolution (Table S1). The model in Fig. 6 was constructed under the assumption that the lever arm behaves as a rigid body in rigor. This assumption is less likely to be valid in other biochemical states, such as relaxation and contraction.

### The N-lobe of RLC is less immobilized than the C-lobe in HMM

Assuming that the motions of the B and E helices of the RLC represent the motion of N- and C-lobes, respectively, we observe that the two lobes behave differently from each other in HMM (Fig. 5, top). While the spectrum of the E helix is similar to that in BSL-RLC-fiber (Fig. 5, bottom), the N-lobe is much more disordered. We hypothesize that this behavior is observed because the N-lobe is adjacent to the flexible C-terminal end of the lever arm.

### Comparison with fluorescence polarization data

To better contextualize our modeling results, we made comparisons with recent fluorescence anisotropy measurements (Romano et al., 2012). In that work, the authors defined a “lever axis” as a vector joining the α-carbons of Cys707 and Lys843 of the 2MYS model. They also defined a “hook axis” as a vector joining the midpoints between Phe836/Ile838 and Met832/Leu834 of the same model. The orientation of the lever arm was characterized by β (the angle between the lever axis and actin), and γ (rotation of the hook axis around the lever axis). Romano et al. (2012) reported a maximum entropy distribution centered at (β, γ) ≈ (105º, 40º). By setting (β, γ) of 2MYS with BSL probes attached to E49C-confidence interval of our method for the E helix, whereas the orientation of the B helix differs from ours by 10.3º. Even though fluorescence anisotropy experiments are versatile in measuring changes of orientation with high temporal resolution, the ME distribution is a low-resolution approximation of the actual angular distribution, while EPR detects this angle with high resolution. so the difference in 10.3° could be attributed to multiple factors.

### Oriented rigor population of myosin with lever arm perpendicular to actin

Previous EPR measurements with monofunctional labels showed at least a 30º lever arm rotation during isometric contraction in scallop muscle (Baker et al., 1998), and at least a 13º lever arm rotation between the rigor and pPDM state (which resembles the pre-power stroke state) in rabbit muscle (Mello and Thomas, 2012). In both of these studies, the lack of stereospecific labeling prevented the accurate measurement of lever arm orientation.

A key conclusion of the present study, that the lever arm occupies a more perpendicular state in the post-power stroke state in rigor muscle than in the conventional lever-arm down model (based primarily on crystal structures or cryo EM involving myosin fragments), is consistent with other structural studies of muscle fibers. EM tomography on highly-ordered insect flight muscle (Schmitz et al., 1996; Chen et al., 2002) shows populations of cross-bridges with more perpendicular lever arm orientation than is predicted by 5H53. Perpendicular conformation of a myosin lever arm was found in two types of cross-bridges, termed “lead bridge” and “reaer bridge.” The lead bridge is a double-headed cross-bridge in a myac layer (longitudinal section containing a single layer of alternating actin and myosin filaments). The myosin molecule that is closer to M-line (M-ward lead bridge head) is fitted by a “lever arm down” model (tilt angle 151° ± 1.1°). The myosin molecule that is closer to the Z-disk however (Z-ward lead bridge head) has a tilt angle of 98° ± 2.3°. The rear bridge is a single-headed cross-bridge with a lever arm tilt of 106° ± 12°. Our EPR-derived model (Fig. 6) is compatible with either of these approximately perpendicular orientations. A characteristic property of the rear bridge EM density is that it disappears under nucleotide addition, e.g. AMPPNP (Schmitz et al., 1996). Addition of 5mM AMPPNP, however, does not change the angular distribution in our case (Fig. S4). Thus, the population of oriented lever arms that we observe might correspond to a structure resembling the Z-ward heads of the lead bridge. In the EM tomography analysis, group averaging was performed over self-similar motifs. This method prevents averaging-out of ordered populations in a sample with a high degree of heterogeneity. Similarly, EPR spectra are highly sensitive to ordered populations, even when they are sparsely populated.

Previous FRET studies on fast skeletal muscle fibers with the A2 ELC isoform are consistent with a perpendicular lever arm orientation in rigor (Guhathakurta et al., 2015), as observed more directly in the present study by EPR. Applying our approach to cardiac muscle fibers might shed additional light on this hypothesis, since cardiac muscle has only the A2 ELC isoform.

## CONCLUSION

In the present study, we used stereospecific labeling of RLC at two distinct sites to determine the orientation of the lever arm of actin-attached myosin in skinned skeletal muscle fibers in rigor (no ATP). Two populations were observed simultaneously, both ordered and disordered, but the only ordered population has a lever arm that is perpendicular to the muscle fiber axis. This orientation, determined in muscle fibers at ambient temperature, differs by 33° from the “lever arm down” orientation determined from cryo-EM on isolated myosin heads bound to actin. Relaxation (addition of ATP in the absence of Ca) produced substantial angular disorder. These results have a profound effect on models of muscle force generation, and future experiments under other physiological conditions have the potential to provide a more direct link between muscle physiology and structural biology.

## AUTHOR CONTRIBUTIONS

All authors participated in the experimental design. YS performed the research and analyzed the data. BPB performed molecular modeling. YS, BPB, ART, and DDT wrote the paper. All authors approved the final version of the manuscript.

## ACKNOWLEDGEMENTS

We thank Malcolm Irving and Luca Fusi for fruitful discussions, Kenneth Taylor for an extensive feedback, Edmund Howard for EPR discussions, Lien Phung and John Rohde for proofreading the manuscript, Margaret Titus for training, and Octavian Cornea for administrative assistance. We used the facilities and expertise of the University of Minnesota Center for Mass Spectrometry and Proteomics for data acquisition and analysis. EPR experiments were performed at the Biophysical Technology Center, University of Minnesota. This study was supported by National Institutes of Health (NIH) grants R01AR032961 and R37AG26160 to DDT. YS was supported by NIH T32AR007612 and by a University of Minnesota Interdisciplinary Doctoral Fellowship.

## SUPPLEMENTAL MATERIAL

**Table S1.** Fitting parameters of the oriented fiber experiments.

|  | B helix (53-57) |  |  |
| --- | --- | --- | --- |
| | $\theta_{NB}(^{\circ})$ | $\sigma(^{\circ})$ | Mole-fraction |
| RLC-fiber rigor (first component) | 4±4 | 9±3 | 0.11±0.02 |
| (second component) | 22±15 | >38 | 0.89±0.02 |
| RLC-fiber relaxation (single component) | 18±14 | >38 | - |
| RLC-HMM-fiber rigor (single component) | 32±14 | >38 | - |
|  | E helix (103-107) |  |  |
| | $\theta_{NB}(^{\circ})$ | $\sigma(^{\circ})$ | Mole-fraction |
| RLC-fiber rigor (first component) | 81±4 | 11±3 | 0.54±0.03 |
| (second component) | 86±18 | >38 | 0.46±0.03 |
| RLC-fiber relaxation (single component) | 51±15 | >38 | - |
| RLC-HMM-fiber rigor (first component) | 81±4 | 11±3 | 0.25±0.03 |
| (second component) | 88±16 | >38 | 0.75±0.03 |

Errors are propagated from 95% confidence intervals imposed on the simulated fit error surface.

**Fig. S1.**
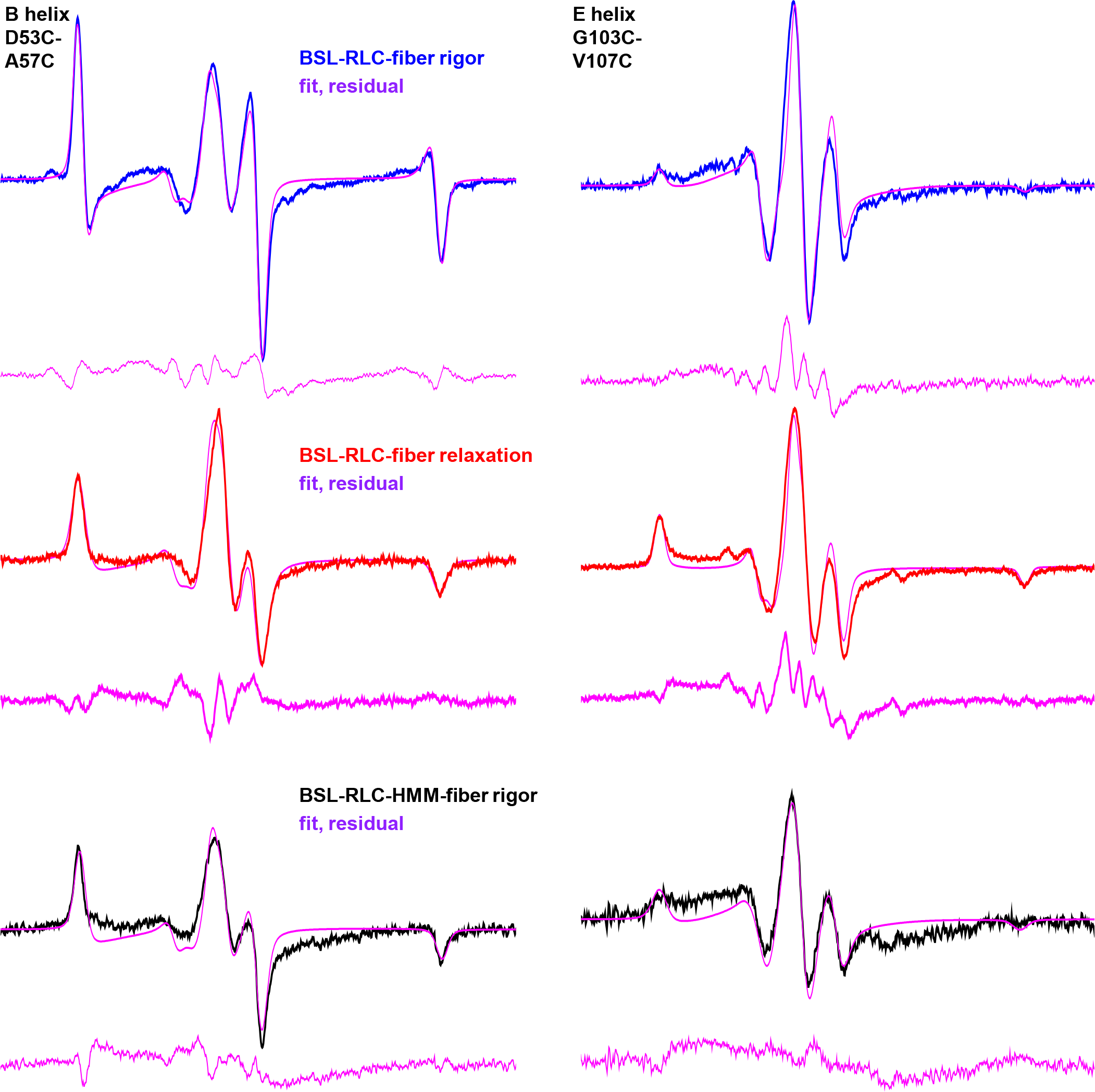
EPR data on BSL-RLC-fiber in rigor (blue), BSL-RLC-fiber in relaxation (red) and BSL-RLC-HMM-fiber in rigor (black). Fits and residuals are in magenta. Static regime was assumed during the fitting. Field sweep is 100 G.

**Fig. S2.**
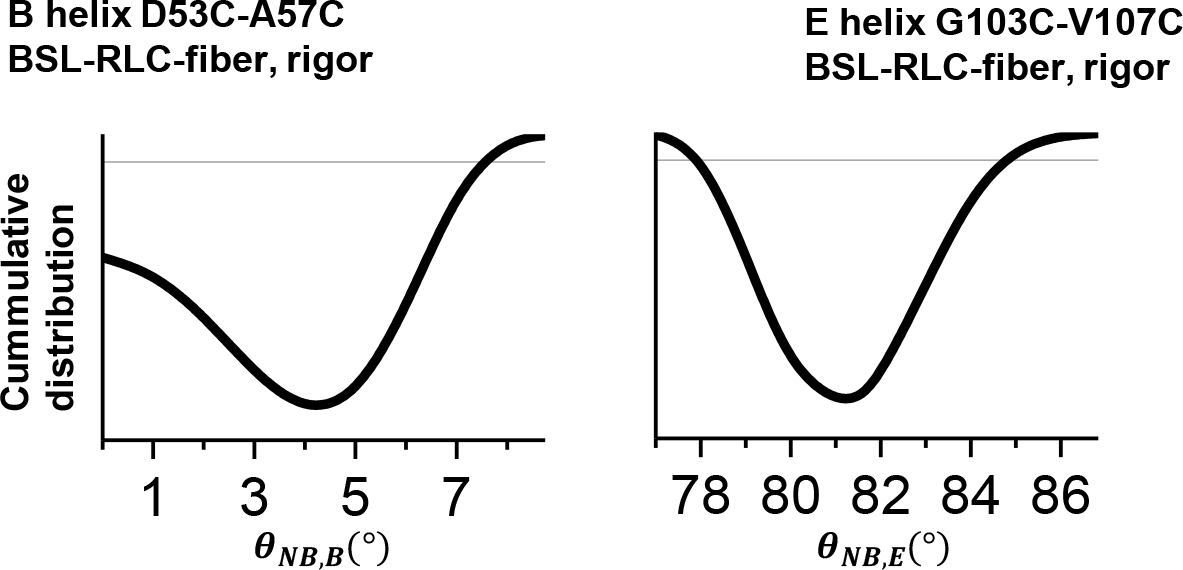
Cumulative distribution function of the F-ratio distribution, comparing the ratio of the residual sum of squares of the fits with the residual sum of squares of the best fit. The 95% intervals are depicted by horizontal lines.

**Fig. S3.**
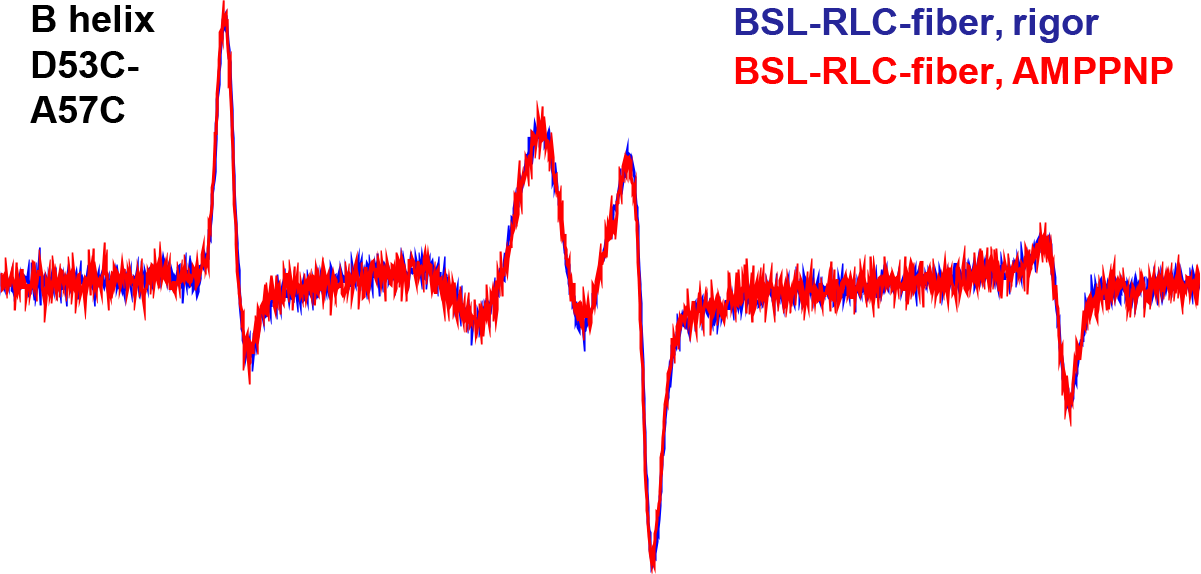
Effect of 5 mM AMPPMP (red) on BSL-RLC-fiber labeled on the B helix in rigor (blue) during EPR. Field sweep is 100 G.

